## Supplementary Tables and Figures for "Multi-dimensionality of tree communities structure host-parasitoid networks and their phylogenetic composition"

***Wang et al.***

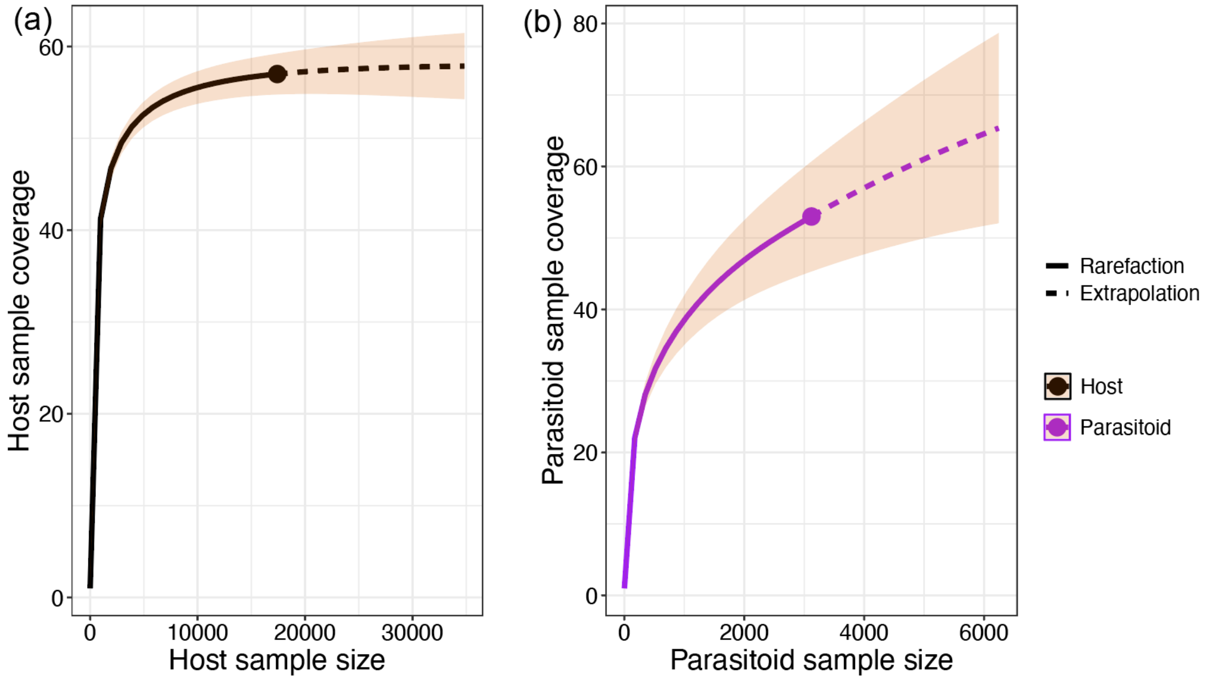


Fig. S1 The sample coverage across different sample sizes for a) hosts and b) parasitoids.

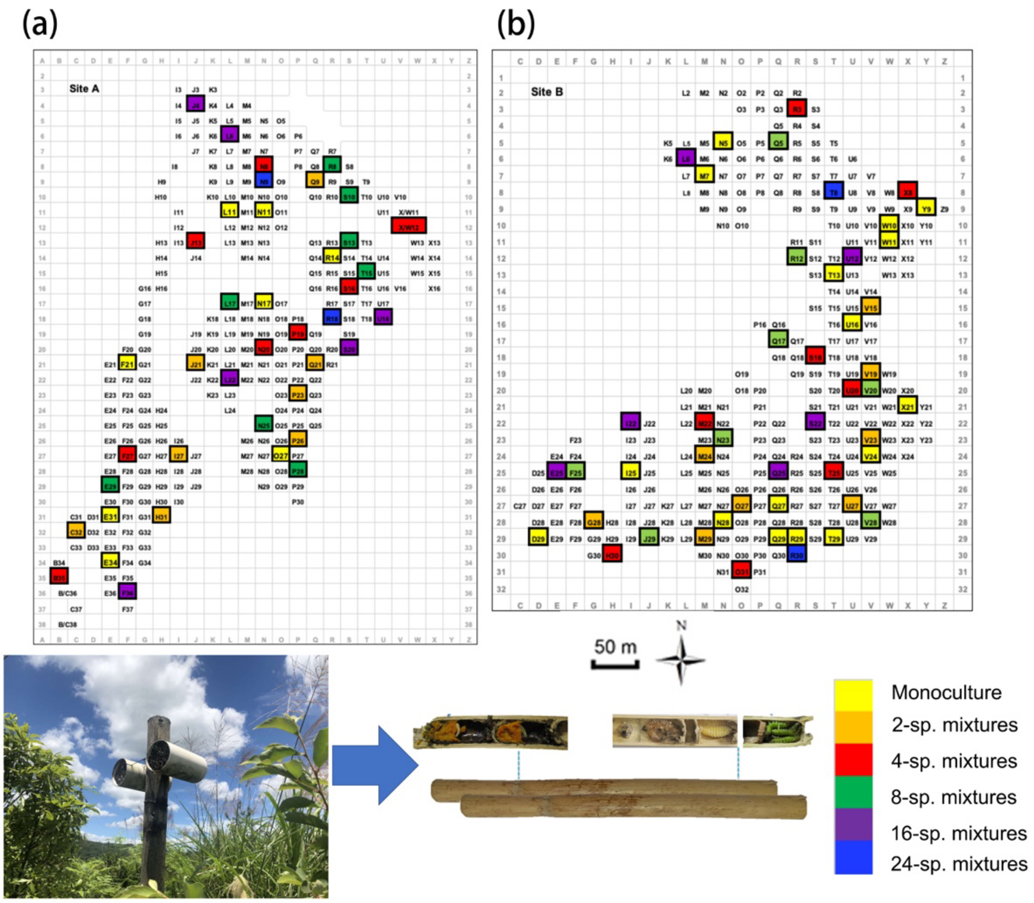


Fig. S2. Overview of the study plot distribution along the two experimental tree diversity sites of BEF-China (a: Site A, b: Site B).Levels of tree species richness indicated by color. Each study plot had a size of 25.8 m x 25.8 m.

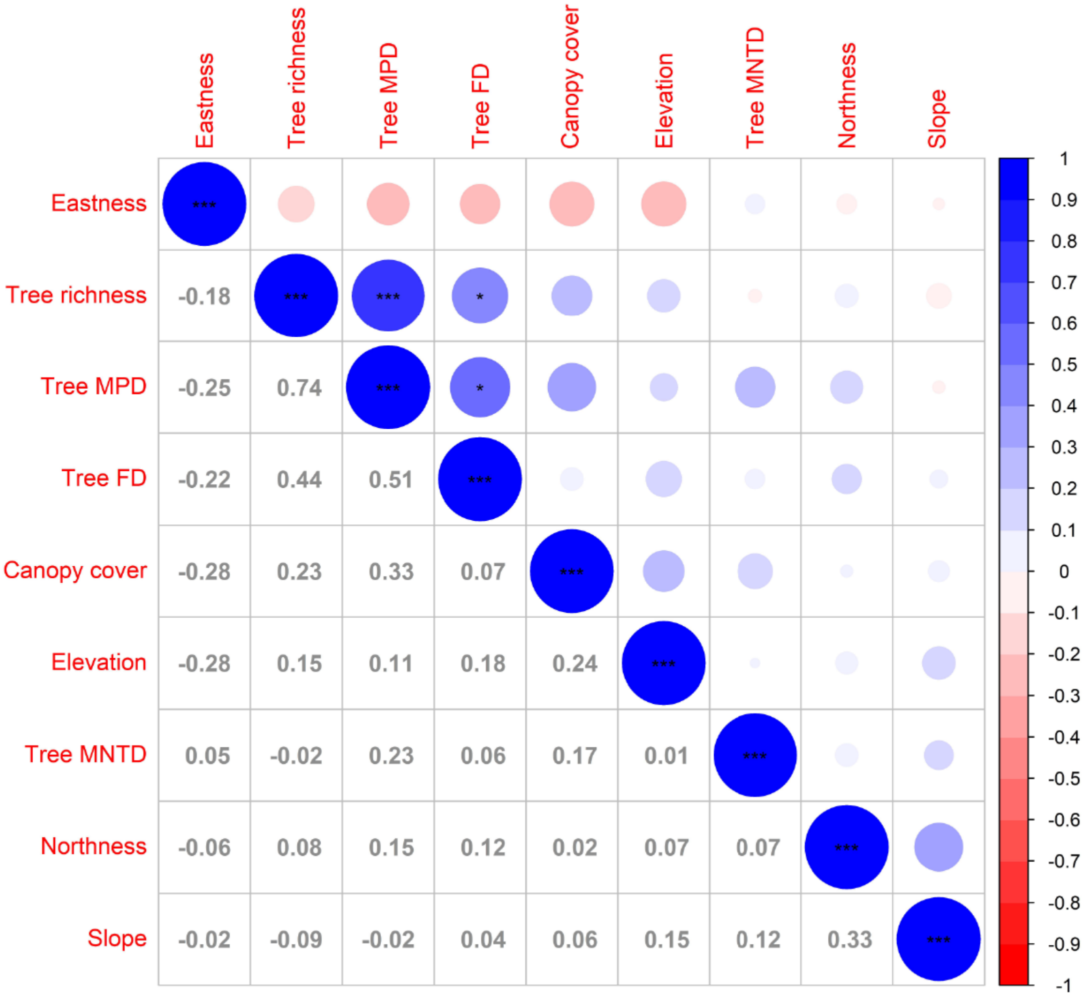


Fig. S3. Correlations among the predictors used in the study. Values and colored circles are Pearson's correlation coefficients r, significances are indicated by asterisks: *** P < 0.001, ** P < 0.01, * P ⩽ 0.05. Tree MPD (tree mean pairwise phylogenetic distance), Tree FD (tree functional diversity expressed as Rao's Q), Tree MNTD (tree mean nearest taxon distance).

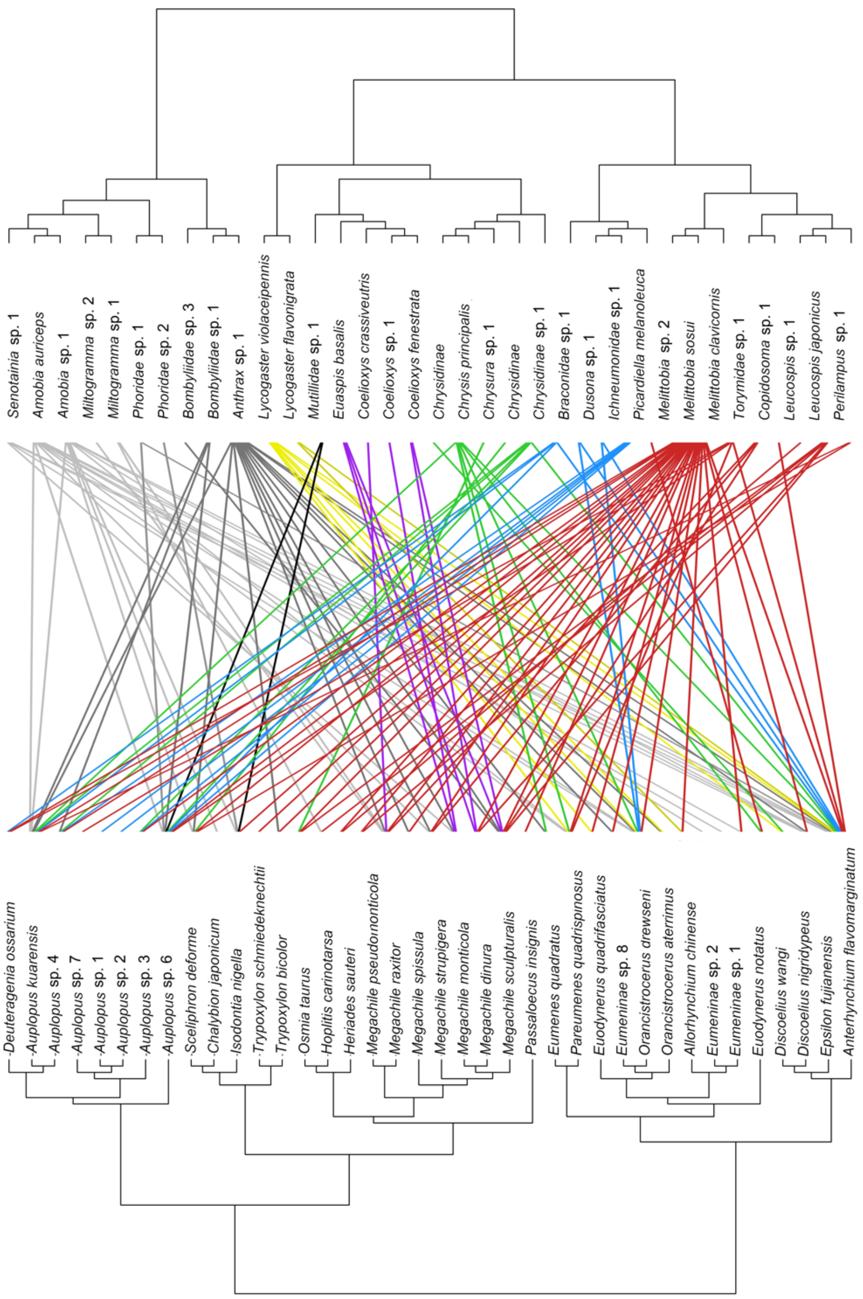


Fig. S4. Dendrogram of phylogenetic congruence for the host species (below) and associated parasitoid species (above) recorded in the study, showing the species name for hosts and parasitoids.

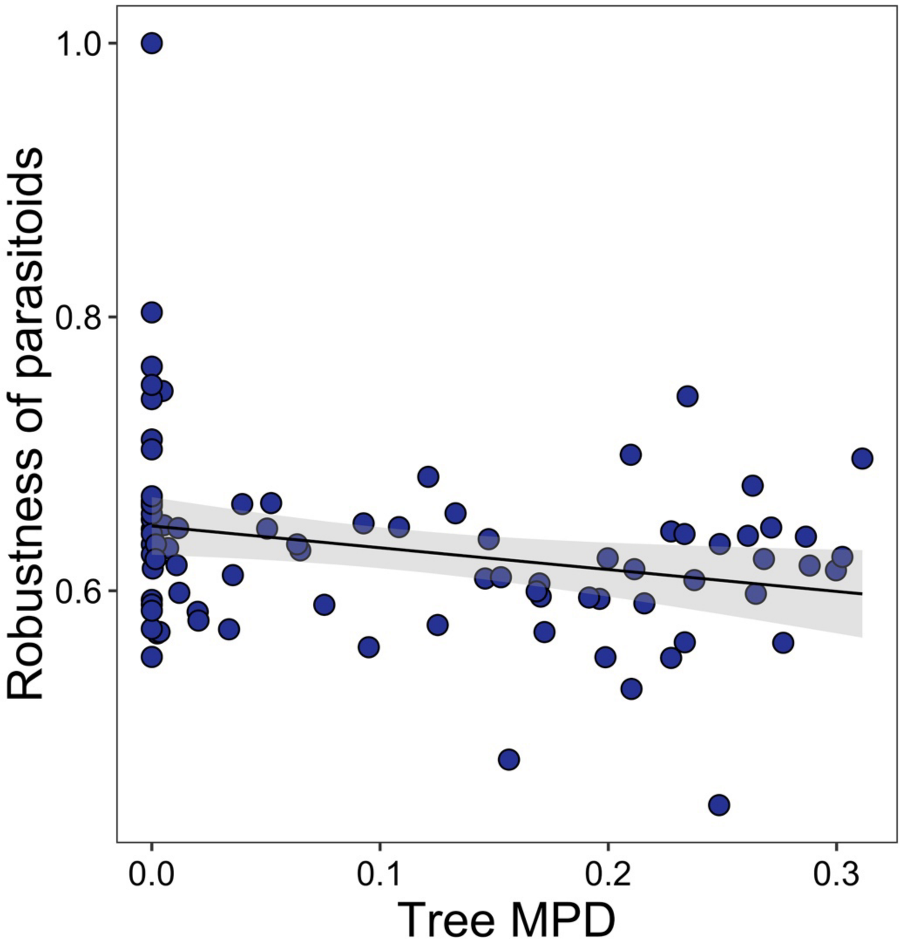


Fig. S5. Community-level relationships of networks between tree MPD and robustness of parasitoids. Values were adjusted for covariates of the final regression model (i.e. partial residuals shown on y-axis). Regression lines (with 95% confidence bands) show significant ($p < 0.05$) relationships.

Table S1. Hymenopteran species composition and abundance across all study plots species used in parafit analysis were marked.

| Family | Species | Abundance | Role | Parafit |
|---|---|---|---|---|
| Colletidae | *Hylaeus* sp.1 | 62 | Host | |
| Megachilidae | *Heriades sauteri* | 201 | Host | √ |
| Megachilidae | *Hoplitis carinotarsa* | 382 | Host | √ |
| Megachilidae | *Megachile strupigera* | 326 | Host | √ |
| Megachilidae | *Megachile monticola* | 18 | Host | √ |
| Megachilidae | *Megachile pseudomonticola* | 10 | Host | √ |
| Megachilidae | *Megachile rixato*r | 131 | Host | √ |
| Megachilidae | *Megachile sculpturalis* | 363 | Host | √ |
| Megachilidae | *Megachile spissula* | 313 | Host | √ |
| Megachilidae | *Megachile dinura* | 7 | Host | √ |
| Megachilidae | *Osmia taurus* | 121 | Host | √ |
| Megachilidae | *Trachusa staabi* | 83 | Host | |
| Vespidae | *Allorhynchium chinense* | 83 | Host | √ |
| Vespidae | *Anterhynchium flavomarginatum* | 8644 | Host | √ |
| Vespidae | *Discoelius nigridypeus* | 15 | Host | √ |
| Vespidae | *Discoelius* sp.1 | 4 | Host | |
| Vespidae | *Discoelius wangi* | 41 | Host | √ |
| Vespidae | *Epsilon fujianensis* | 483 | Host | √ |
| Vespidae | *Eumenes quadratus* | 17 | Host | √ |
| Vespidae | *Eumenes sillicus* | 4 | Host | |
| Vespidae | *Eumenes* sp.1 | 3 | Host | |
| Vespidae | Eumeninae sp.1 | 2 | Host | √ |
| Vespidae | Eumeninae sp.2 | 19 | Host | √ |
| Vespidae | Eumeninae sp.3 | 3 | Host | |
| Vespidae | Eumeninae sp.4 | 11 | Host | |
| Vespidae | Eumeninae sp.5 | 1 | Host | |
| Vespidae | Eumeninae sp.8 | 8 | Host | √ |
| Vespidae | *Euodynerus notatus* | 54 | Host | √ |
| Vespidae | *Euodynerus quadrifasciatus* | 49 | Host | √ |
| Vespidae | *Orancistrocerus aterrimus* | 34 | Host | √ |
| Vespidae | *Orancistrocerus drewseni* | 367 | Host | √ |
| Vespidae | *Pararrhynchium ornatum* | 16 | Host | |
| Vespidae | *Pareumenes quadrispinosus* | 389 | Host | √ |
| Sphecidae | *Chalybion japonicum* | 53 | Host | √ |
| Sphecidae | *Isodontia nigella* | 109 | Host | √ |
| Sphecidae | *Sceliphron deforme* | 90 | Host | √ |
| Crabronidae | *Passaloecus insignis* | 27 | Host | √ |
| Crabronidae | *Pison atripenne* | 31 | Host | |
| Crabronidae | Crabronidae sp.1 | 14 | Host | |

| | | | | |
|---|---|---|---|---|
| Crabronidae | *Trypoxylon bicolor* | 91 | Host | √ |
| Crabronidae | *Trypoxylon* sp. | 26 | Host | √ |
| Pompilidae | *Auplopus kuarensis* | 371 | Host | √ |
| Pompilidae | *Auplopus carbonarius* | 141 | Host | |
| Pompilidae | *Auplopus* sp.1 | 1 | Host | √ |
| Pompilidae | *Auplopus* sp.2 | 29 | Host | √ |
| Pompilidae | *Auplopus* sp.3 | 13 | Host | √ |
| Pompilidae | *Auplopus* sp.4 | 66 | Host | √ |
| Pompilidae | *Auplopus* sp.5 | 1 | Host | |
| Pompilidae | *Auplopus* sp.6 | 92 | Host | √ |
| Pompilidae | *Auplopus* sp.7 | 3 | Host | √ |
| Pompilidae | *Auplopus* sp.9 | 4 | Host | |
| Pompilidae | *Auplopus* sp.10 | 2 | Host | |
| Pompilidae | *Auplopus* sp.11 | 10 | Host | |
| Pompilidae | *Auplopus* sp.12 | 5 | Host | |
| Pompilidae | *Deuteragenia ossarium* | 110 | Host | √ |
| Pompilidae | Pompilidae sp.1 | 3 | Host | |
| Braconidae | Braconidae sp.1 | 32 | parasitoid | √ |
| Eulophidae | *Chaenotetrastichus semiflavus* | 4 | parasitoid | |
| Eulophidae | Eulophidae sp.1 | 1 | parasitoid | |
| Eulophidae | *Kocourekia* sp.1 | 11 | parasitoid | |
| Eulophidae | *Melittobia australica* | 1 | parasitoid | |
| Eulophidae | *Melittobia clavicornis* | 42 | parasitoid | √ |
| Eulophidae | *Melittobia sosui* | 444 | parasitoid | √ |
| Eulophidae | *Melittobia* sp.1 | 1 | parasitoid | |
| Eulophidae | *Melittobia* sp.2 | 2 | parasitoid | √ |
| Chrysididae | Chrysidinae sp.1 | 76 | parasitoid | √ |
| Chrysididae | Chrysidinae sp.2 | 4 | parasitoid | |
| Chrysididae | Chrysidinae sp.3 | 2 | parasitoid | |
| Chrysididae | Chrysidinae sp.4 | 2 | parasitoid | √ |
| Chrysididae | Chrysidinae sp.5, | 6 | parasitoid | √ |
| Chrysididae | *Chrysis principalis* | 1244 | parasitoid | √ |
| Chrysididae | *Chrysura* sp.1 | 9 | parasitoid | √ |
| Encyrtidae | *Copidosoma* sp.1 | 1 | parasitoid | √ |
| Ichneumonidae | *Dusona* sp.1 | 35 | parasitoid | √ |
| Ichneumonidae | Ichneumonidae sp.1 | 22 | parasitoid | √ |
| Ichneumonidae | Ichneumonidae sp.5 | 4 | parasitoid | |
| Ichneumonidae | Ichneumonidae sp.6 | 2 | parasitoid | |
| Ichneumonidae | Ichneumonidae sp.7 | 1 | parasitoid | |
| Ichneumonidae | *Picardiella melanoleuca* | 90 | parasitoid | √ |
| Leucospidae | *Leucospis japonicas* | 27 | parasitoid | √ |
| Leucospidae | *Leucospis* sp.1 | 6 | parasitoid | √ |
| Trigonalyidae | *Lycogaster flavonigrata* | 4 | parasitoid | √ |

| | | | | |
|---|---|---|---|---|
| Trigonalyidae | *Lycogaster violaceipennis* | 597 | parasitoid | √ |
| Mutillidae | Mutillidae sp.1 | 8 | parasitoid | √ |
| Mutillidae | Mutillidae sp.2 | 1 | parasitoid | |
| Perilampidae | Perilampidae sp.1 | 3 | parasitoid | √ |
| Torymidae | Torymidae sp.1 | 28 | parasitoid | √ |
| Megachilidae | *Coelioxys crassiventris* | 75 | parasitoid | √ |
| Megachilidae | *Coelioxys fenestrate* | 52 | parasitoid | √ |
| Megachilidae | *Coelioxys* sp.1 | 1 | parasitoid | √ |
| Megachilidae | *Euaspis basalis* | 32 | parasitoid | √ |
| Megachilidae | *Euaspis Polynesia* | 4 | parasitoid | |
| Gasteruptiidae | *Gasteruption corniculigerum* | 14 | parasitoid | |
| Bombyliidae | *Anthrax* sp.1 | 127 | parasitoid | √ |
| Bombyliidae | Bombyliidae sp.1 | 6 | parasitoid | √ |
| Bombyliidae | Bombyliidae sp.2 | 1 | parasitoid | |
| Bombyliidae | Bombyliidae sp.3 | 1 | parasitoid | √ |
| Phoridae | Phoridae sp.1 | 7 | parasitoid | √ |
| Phoridae | Phoridae sp.2 | 1 | parasitoid | √ |
| Sarcophagidae | *Amobia auriceps* | 855 | parasitoid | √ |
| Sarcophagidae | *Amobia* sp.1 | 125 | parasitoid | √ |
| Sarcophagidae | *Miltogramma* sp.1 | 4 | parasitoid | √ |
| Sarcophagidae | *Miltogramma* sp.2 | 1 | parasitoid | √ |
| Sarcophagidae | *Miltogramma* sp.3 | 1 | parasitoid | |
| Sarcophagidae | *Senotainia* sp.1 | 26 | parasitoid | √ |

Table S2. Environmental correlates of host species community dissimilarity (NMDS on Morisita-Horn dissimilarity) across the study plots. Correlation coefficients, explained variation ($r^2$) and probabilities (based on 1000 permutations) for the relationships between environmental variables and the NMDS axis scores. Significant P-values are indicated in bold.

| Vectors | Host species composition | | | |
|---|---|---|---|---|
| | NMDS1 | NMDS2 | $r^2$ | Pr(>r) |
| Tree phylogeny NMDS1 | -0.9674 | 0.25327 | 0.0359 | 0.225 |
| Tree phylogeny NMDS2 | 0.26373 | -0.9646 | 0.1517 | **0.003** |
| Tree composition NMDS1 | 0.62106 | -0.78377 | 0.3298 | **0.001** |
| Tree composition NMDS2 | 0.16588 | 0.98615 | 0.0127 | 0.604 |
| Canopy cover | -0.99998 | -0.00593 | 0.5095 | **0.001** |
| Tree species richness | -0.90074 | -0.43436 | 0.0798 | 0.035 |
| Elevation | -0.65845 | 0.75262 | 0.13 | **0.005** |
| Eastness | 0.79555 | 0.60589 | 0.0586 | 0.079 |
| Northness | -0.07337 | -0.99731 | 0.0171 | 0.49 |
| Slope | -0.13253 | -0.99118 | 0.0836 | 0.031 |
| Tree FD (Rao's Q) | -0.97061 | 0.24067 | 0.0549 | 0.094 |
| Tree MPD | -0.93721 | -0.34878 | 0.1439 | **0.005** |
| Parasitoid phylogeny NMDS1 | 0.93994 | -0.34133 | 0.187 | **0.001** |
| Parasitoid phylogeny NMDS2 | -0.32919 | 0.94426 | 0.0183 | 0.462 |
| Parasitoid compositionNMDS1 | -0.94691 | 0.3215 | 0.1881 | **0.001** |
| Parasitoid compositionNMDS2 | 0.9524 | -0.30484 | 0.106 | **0.014** |
| stress=0.23 | | | | |

Table S3. Environmental correlates of parasitoid species community dissimilarity (NMDS on Morisita-Horn dissimilarity) across the study plots. Correlation coefficients, explained variation ($r^2$) and probabilities (based on 1000 permutations) for the relationships between environmental variables and the NMDS axis scores. Significant P-values are indicated in bold.

| Vectors | Parasitoid species composition | | | |
|---|---|---|---|---|
| | NMDS1 | NMDS2 | $r^2$ | Pr(>r) |
| Tree phylogeny NMDS1 | 0.19179 | -0.98144 | 0.0217 | 0.422 |
| Tree phylogeny NMDS2 | -0.47649 | 0.87918 | 0.0518 | 0.12 |
| Tree composition NMDS1 | -0.55799 | 0.82985 | 0.2263 | **0.001** |
| Tree composition NMDS2 | -0.50111 | 0.86538 | 0.0201 | 0.418 |
| Canopy cover | 0.8741 | -0.48575 | 0.2736 | **0.001** |
| Tree species richness | 0.98384 | 0.17905 | 0.0524 | 0.122 |
| Elevation | 0.55299 | -0.83319 | 0.1263 | **0.007** |
| Eastness | -0.59888 | 0.80084 | 0.0717 | 0.045 |
| Northness | -0.81065 | 0.58554 | 0.0045 | 0.837 |
| Slope | 0.95482 | 0.29719 | 0.0176 | 0.507 |
| Tree FD (Rao's Q) | 0.88491 | -0.46576 | 0.0921 | **0.019** |
| Tree MPD | 0.99591 | 0.09035 | 0.0972 | **0.021** |
| Host phylogeny NMDS1 | -0.93007 | 0.36738 | 0.1688 | **0.016** |
| Host phylogeny NMDS2 | -0.93952 | -0.3425 | 0.0834 | **0.027** |
| Host compositionNMDS1 | -0.96573 | 0.25953 | 0.3706 | **0.001** |
| Host compositionNMDS2 | 0.52611 | -0.85042 | 0.0458 | 0.169 |
| stress=0.23 | | | | |

Table S4. Environmental correlates of host phylogenetic community dissimilarity (NMDS on Morisita-Horn dissimilarity) across the study plots. Correlation coefficients, explained variation ($r^2$) and probabilities (based on 1000 permutations) for the relationships between environmental variables and the NMDS axis scores. Significant P-values are indicated in bold.

| Vectors | Host phylogenetic composition | | | |
|---|---|---|---|---|
| | NMDS1 | NMDS2 | $r^2$ | Pr(>r) |
| Tree phylogeny NMDS1 | 0.19179 | -0.98144 | 0.0217 | 0.386 |
| Tree phylogeny NMDS2 | -0.47649 | 0.87918 | 0.0518 | 0.128 |
| Tree composition NMDS1 | -0.55799 | 0.82985 | 0.2263 | **0.001** |
| Tree composition NMDS2 | -0.50111 | 0.86538 | 0.0201 | 0.433 |
| Canopy cover | 0.8741 | -0.48575 | 0.2736 | **0.001** |
| Tree species richness | 0.98384 | 0.17905 | 0.0524 | 0.100 |
| Elevation | 0.55299 | -0.83319 | 0.1263 | **0.001** |
| Eastness | -0.59888 | 0.80084 | 0.0717 | **0.04** |
| Northness | -0.81065 | 0.58554 | 0.0045 | 0.821 |
| Slope | 0.95482 | 0.29719 | 0.0176 | 0.507 |
| Tree FD (Rao's Q) | 0.88491 | -0.46576 | 0.0921 | **0.021** |
| Tree MPD | 0.99591 | 0.09035 | 0.0972 | **0.013** |
| Parasitoid phylogeny NMDS1 | -0.49168 | 0.87078 | 0.4286 | **0.001** |
| Parasitoid phylogeny NMDS2 | 0.61112 | -0.79154 | 0.0714 | 0.058 |
| Parasitoid compositionNMDS1 | 0.42314 | -0.90606 | 0.7623 | **0.001** |
| Parasitoid compositionNMDS2 | -0.95942 | -0.28198 | 0.2125 | **0.001** |
| stress=0.24 | | | | |

Table S5. Environmental correlates of parasitoid phylogenetic community dissimilarity (NMDS on Morisita-Horn dissimilarity) across the study plots. Correlation coefficients, explained variation ($r^2$) and probabilities (based on 1000 permutations) for the relationships between environmental variables and the NMDS axis scores. Significant P-values are indicated in bold.

| Vectors | Parasitoid phylogenetic composition | | | |
|---|---|---|---|---|
| | NMDS1 | NMDS2 | $r^2$ | Pr(>r) |
| Tree phylogeny NMDS1 | -0.97914 | 0.20321 | 0.0305 | 0.274 |
| Tree phylogeny NMDS2 | 0.99965 | -0.02634 | 0.0889 | **0.024** |
| Tree composition NMDS1 | 0.82944 | -0.5586 | 0.2794 | **0.001** |
| Tree composition NMDS2 | 0.46615 | -0.8847 | 0.0789 | **0.031** |
| Canopy cover | -0.97111 | -0.23861 | 0.1497 | **0.004** |
| Tree species richness | -0.99834 | 0.05759 | 0.0594 | 0.094 |
| Elevation | -0.97095 | -0.23928 | 0.2242 | **0.001** |
| Eastness | 0.95102 | -0.30912 | 0.1682 | **0.001** |
| Northness | -0.27806 | -0.96056 | 0.0274 | 0.340 |
| Slope | -0.37915 | 0.92534 | 0.001 | 0.959 |
| Tree FD (Rao's Q) | -0.88976 | 0.45642 | 0.0819 | **0.031** |
| Tree MPD | -0.98331 | 0.18195 | 0.0363 | 0.223 |
| Host phylogeny NMDS1 | -0.2796 | -0.96012 | 0.0105 | 0.584 |
| Host phylogeny NMDS2 | -0.37192 | 0.92826 | 0.0022 | 0.914 |
| Host compositionNMDS1 | 0.92516 | 0.37957 | 0.1225 | **0.008** |
| Host compositionNMDS2 | -0.9934 | 0.11469 | 0.0485 | 0.138 |
| stress=0.20 | | | | |

Table S6. Results of permutational multivariate analysis of variance (PERMANOVA) between host species composition and environmental variables.

| | DF | Sum of Sqs | $R^2$ | F | P |
|---|---|---|---|---|---|
| Canopy cover | 1 | 1.88 | 0.15 | 15.88 | 0.001 |
| Tree species richness | 1 | 0.18 | 0.01 | 1.53 | 0.131 |
| Elevation | 1 | 0.37 | 0.03 | 3.09 | 0.020 |
| Eastness | 1 | 0.10 | 0.01 | 0.87 | 0.502 |
| Northness | 1 | 0.10 | 0.01 | 0.85 | 0.490 |
| Slope | 1 | 0.32 | 0.03 | 2.74 | 0.016 |
| Tree FD (Rao's Q) | 1 | 0.10 | 0.01 | 0.87 | 0.500 |
| Tree MPD | 1 | 0.10 | 0.01 | 0.85 | 0.493 |
| Residual | 76 | 9.00 | 0.74 | | |
| Total | 84 | 12.16 | 1.00 | | |

Table S7. Results of permutational multivariate analysis of variance (PERMANOVA) between parasitoid species composition and environmental variables.

| | DF | Sum of Sqs | $R^2$ | F | P |
|---|---|---|---|---|---|
| Canopy cover | 1 | 1.14 | 0.08 | 7.72 | 0.001 |
| Tree species richness | 1 | 0.21 | 0.01 | 1.41 | 0.173 |
| Elevation | 1 | 0.40 | 0.03 | 2.71 | 0.015 |
| Eastness | 1 | 0.24 | 0.02 | 1.62 | 0.127 |
| Northness | 1 | 0.21 | 0.02 | 1.41 | 0.170 |
| Slope | 1 | 0.17 | 0.01 | 1.17 | 0.296 |
| Tree FD (Rao's Q) | 1 | 0.19 | 0.01 | 1.30 | 0.252 |
| Tree MPD | 1 | 0.12 | 0.01 | 0.79 | 0.587 |
| Residual | 76 | 11.19 | 0.81 | | |
| Total | 84 | 13.87 | 1.00 | | |

Table S8. Results of permutational multivariate analysis of variance (PERMANOVA) between host phylogenetic composition and environmental variables.

| | DF | Sum of Sqs | $R^2$ | F | P |
|---|---|---|---|---|---|
| Canopy cover | 1 | 0.00 | 0.01 | 0.90 | 0.385 |
| Tree species richness | 1 | 0.00 | 0.01 | 0.63 | 0.467 |
| Elevation | 1 | 0.00 | 0.00 | 0.31 | 0.666 |
| Eastness | 1 | 0.01 | 0.01 | 1.22 | 0.280 |
| Northness | 1 | 0.00 | 0.00 | 0.22 | 0.768 |
| Slope | 1 | 0.03 | 0.07 | 5.83 | 0.019 |
| Tree FD (Rao’s Q) | 1 | 0.00 | 0.01 | 0.53 | 0.507 |
| Tree MPD | 1 | 0.00 | 0.00 | 0.17 | 0.822 |
| Residual | 75 | 0.37 | 0.88 | | |
| Total | 83 | 0.42 | 1.00 | | |

Table S9. Results of permutational multivariate analysis of variance (PERMANOVA) between parasitoid phylogenetic composition and environmental variables.

| | DF | Sum of Sqs | $R^2$ | F | P |
|---|---|---|---|---|---|
| Canopy cover | 1 | 0.01 | 0.04 | 3.91 | 0.013 |
| Tree species richness | 1 | 0.00 | 0.01 | 0.97 | 0.359 |
| Elevation | 1 | 0.00 | 0.02 | 1.96 | 0.116 |
| Eastness | 1 | 0.00 | 0.02 | 2.18 | 0.096 |
| Northness | 1 | 0.00 | 0.01 | 0.76 | 0.483 |
| Slope | 1 | 0.00 | 0.02 | 1.99 | 0.116 |
| Tree FD (Rao's Q) | 1 | 0.00 | 0.01 | 0.86 | 0.393 |
| Tree MPD | 1 | 0.00 | 0.00 | 0.42 | 0.732 |
| Residual | 75 | 0.12 | 0.85 | | |
| Total | 83 | 0.14 | 1.00 | | |

Table S9. Summary results of Mantel test between community composition of trees, hosts and parasitoids.

| | Species composition | | |
|---|---|---|---|
| | Tree & Host | Tree & Parasitoid | Host & Parasitoid |
| Mantel r | 0.13 | 0.14 | 0.65 |
| p | 0.001 | 0.001 | 0.001 |

| | Phylogenetic composition | | |
|---|---|---|---|
| | Tree & Host | Tree & Parasitoid | Host & Parasitoid |
| Mantel r | 0.07 | 0.14 | 0.06 |
| p | 0.07 | 0.002 | 0.139 |

Table S11. Summary results of alternative linear models for parasitoid generality, host vulnerability, robustness, linkage density, connectance, and interaction evenness of community-level host-parasitoid network indices across values of tree phylogenetic diversity (MPD). Standardized parameter estimates (with standard errors, t and P values) are shown for the variables retained in the minimal models.

| | | Est. | SE | t | P |
|---|---|---|---|---|---|
| Parasitoid generality | Intercept | 0.176 | 0.016 | 10.96 | <0.001 |
| | Canopy cover | 0.033 | 0.016 | 2.03 | 0.046 |
| Host vulnerability | Intercept | 2.878 | 0.08 | 35.79 | <0.001 |
| | Tree species richness: Site A | 0.072 | 0.13 | 0.57 | 0.569 |
| | Tree species richness: Site B | 0.256 | 0.11 | 2.43 | 0.017 |
| Robustness of parasitoids | Intercept | 0.631 | 0.01 | 83.44 | <0.001 |
| | Tree species richness: Site A | -0.023 | 0.01 | -1.95 | 0.055 |
| | Tree species richness: Site B | -0.013 | 0.01 | -1.34 | 0.184 |
| Linkage density | Intercept | 2.039 | 0.04 | 50.98 | <0.001 |
| | Elevation | -0.066 | 0.04 | -1.62 | 0.110 |
| | Tree species richness: Site A | 0.091 | 0.06 | 1.42 | 0.160 |
| | Tree species richness: Site B | 0.128 | 0.05 | 2.44 | 0.017 |
| Interaction evenness | Intercept | 0.511 | 0.009 | 59.12 | 0.025 |
| | Canopy cover | -0.037 | 0.007 | -5.06 | <0.001 |
| | Eastness | -0.018 | 0.007 | -2.50 | 0.015 |
